# BayesKAT: Bayesian Optimal Kernel-based Test for genetic association studies reveals joint genetic effects in complex diseases

**DOI:** 10.1101/2023.10.18.562824

**Authors:** Sikta Das Adhikari, Yuehua Cui, Jianrong Wang

## Abstract

GWAS methods have identified individual SNPs significantly associated with specific phenotypes. Nonetheless, many complex diseases are polygenic and are controlled by multiple genetic variants that are usually non-linearly dependent. These genetic variants are marginally less effective and remain undetected in GWAS analysis. Kernel-based tests (KBT), which evaluate the joint effect of a group of genetic variants, are therefore critical for complex disease analysis. However, choosing different kernel functions in KBT can significantly influence the type I error control and power, and selecting the optimal kernel remains a statistically challenging task. A few existing methods suffer from inflated type 1 errors, limited scalability, inferior power, or issues of ambiguous conclusions. Here, we present a new Bayesian framework, BayesKAT(https://github.com/wangjr03/BayesKAT), which overcomes these kernel specification issues by selecting the optimal composite kernel adaptively from the data while testing genetic associations simultaneously. Furthermore, BayesKAT implements a scalable computational strategy to boost its applicability, especially for high-dimensional cases where other methods become less effective. Based on a series of performance comparisons using both simulated and real large-scale genetics data, BayesKAT outperforms the available methods in detecting complex group-level associations and controlling type I errors simultaneously. Applied on a variety of groups of functionally related genetic variants based on biological pathways, co-expression gene modules, and protein complexes, BayesKAT deciphers the complex genetic basis and provides mechanistic insights into human diseases.

## INTRODUCTION

Deciphering the genetic basis of complex traits, such as the Alzheimer’s disease and autoimmune diseases, plays pivotal roles in functional genomics and precision medicine (1), (2). Based on the advancement in high-throughput sequencing techniques, specific associated genetic variants, e.g., single-nucleotide polymorphisms (SNPs), have been identified for a large panel of phenotypes using Genome-wide Association Studies (GWAS) (3). However, traditional GWAS approaches treat SNPs independently and can only discover individual SNPs that have strong marginal statistical associations with the phenotype of interest. It is well documented that many complex diseases and phenotypes are often associated with multiple genetic variants (4), (5), (6), (7) where an individual variant itself might be weakly associated with the phenotype. In contrast, groups of such SNPs may jointly contribute to the phenotype, potentially mediated via their cooperative participation in important biological processes or pathways (8), (9). Therefore, the traditional GWAS framework of testing individual SNPs separately without considering the correlation structures and the potential interactions among SNPs may not capture the group-wise joint SNP effects. Separate testings of SNP associations by traditional GWAS approaches are also limited to reveal the underlying biological mechanisms of complex phenotypes. Alternative approaches based on multivariate regression significantly suffer from the large degrees of freedom in genome-wide association tests and can substantially lose the statistical power (10).

To overcome this critical challenge, kernel-based testing (KBT) framework has been introduced to test group-wise joint SNP effects (10), (11), (12), (13), (14), (15), (16). By incorporating a kernel function to measure the similarity among genetic variants and compare with the phenotype similarities, the KBT framework simultaneously models the joint effects of multiple genetic variants within a group. Wu et al. 2011 (12) first proposed the widely-used sequence kernel association test (SKAT) model to test rare-variant associations based on sequencing data. As a supervised, versatile and computationally streamlined regression approach, SKAT accesses the associations between genetic variants within a specific region and the trait, which can be either continuous or dichotomous. Covariates are systematically also accounted for in the SKAT framework. As the outputs from the SKAT model, p-values of the statistical associations are generated, facilitating straightforward interpretations of the findings. An R package (17) has been developed for implementing different kinds of kernel-based testing models, including SKAT.

To enable novel discoveries of the genetic basis underlying complex diseases, maximizing statistical power in genome-wide association tests while effectively controlling type 1 errors is strongly desired in the design of efficient models and algorithms. Under the KBT framework, statistical power heavily depends on the specific choice of kernel functions (16), (18), (19), (20). However, the existing KBT models, including SKAT, require the kernel function to be specified a priori. Because the true functional relationship between the genetic variants and phenotypes is usually unknown in practice, selecting the ideal kernel function in advance for the KBT model, one that maximizes statistical power without increasing the type 1 error rate, poses statistical and computational challenges. One common approach that has been used is to repeat the KBT procedures based on different choices of kernels and then select the one resulting in the minimum p-value, which has been discussed by multiple studies (18), (21). The major problem of this straightforward approach is the inflated type 1 error. Although data-dependent permutation or perturbation methods (18) can help ease the problem, they are not computationally scalable, especially when applied to high-dimensional datasets in large-scale genomic studies. An alternative approach is to use an equal-weighted average of multiple candidate kernels to form an averaged composite kernel (18), which performs better than the worst performing candidate kernel but does not usually achieve the performance of the optimal kernel function. Tests based on the average kernel approach may lead to inconsistent or incorrect conclusions in applications, as we will demonstrate below. He et al. 2018 (21) proposed a maximum kernel test model based on the U statistic, i.e. the mKU model, which claims to achieve the statistical power as close as to the best candidate kernel in high-dimensional settings under certain distributional assumptions. However, the specific distributional assumptions may not hold in practice and hence lead to inflated p-values, which will also be discussed in this study.

To further illustrate the significance and difficulty of choosing appropriate kernels in genetic association testings, Figure 1(A) demonstrates an example based on the genotype data for the trait of whole brain volume collected from the ADNI project for the Alzheimer’s Disease Neuroimaging Initiative ^1^ (https://adni.loni.usc.edu/) (22). The group of genetic variants located in genes belonging to the caffeine metabolism pathway (23), (24), (25) are included into the kernel-based testing model to test the hypothesis that whether the caffeine metabolism pathway is associated with the whole brain volume phenotype. Using different kernel functions, the SKAT model leads to inconsistent conclusions. For instance, based on Quadratic kernel, the SKAT model rejects the null hypothesis (p-value*<*0.05), while the use of Gaussian kernel or IBS kernel in the model does not lead to any rejection of the null hypothesis (Figure 1(A)). On the other hand, using the equal-weighted average composite kernel, the SKAT model tends to reject the null hypothesis (p-value= 0.047). Since there is no clear mechanistic link between the caffeine metabolism pathway and whole brain volume, the rejection of the null hypothesis based on the Quadratic and the average composite kernels is likely a false discovery. To further quantify this issue, in Figure 1(B), a cohort of total 500 replicate synthetic datasets based on the same covariates and genotype data as in Figure 1(A) is created, where the phenotype variables are generated by a Quadratic function *h*^*′*^(.). Applying the SKAT test based on different kernel functions on the synthetic datasets, inconsistent testing results appear to be a persistent problem. Although the Quadratic kernel function leads to the correct hypothesis testing result as expected, the average composite kernel usually leads to incorrect conclusions (Figure 1(B)). As shown in the barplot of Figure 1(B), using different kernels (Linear, Quadratic, Gaussian and average composite kernels) across the 500 synthetic datasets, inconsistent results are observed and the overall fractions of correct testing results are very low. Hence, the inconsistent conclusions based on different kernels suggest the fundamental need of developing a systematic data-adaptive approach of selecting appropriate kernel functions for KBT models in genome-wide association tests.

**Figure 1.**
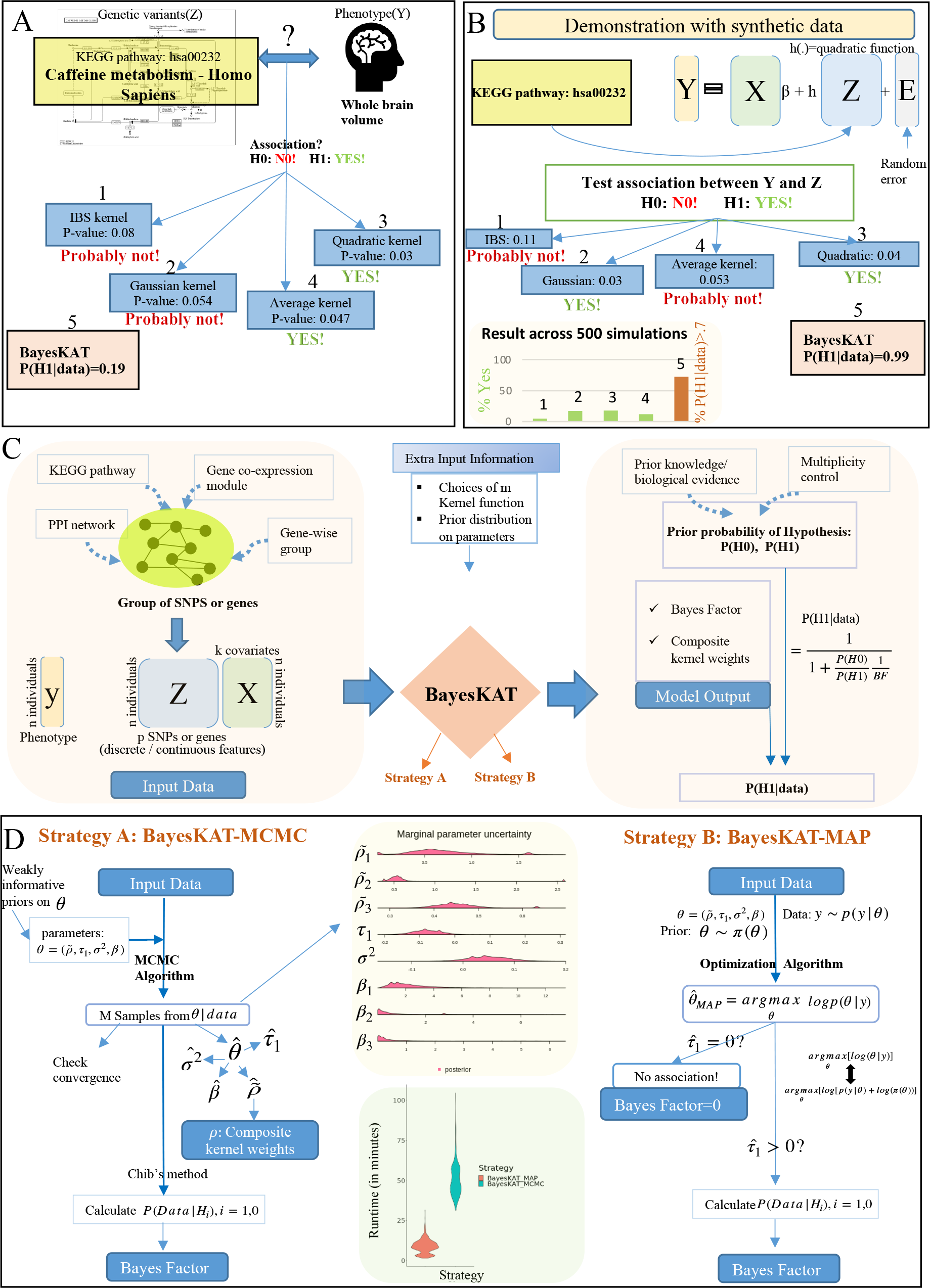
Overview of the significance and model design for BayesKAT. (A) Real-world example of association tests with inconsistent results depending on specific kernels. To test the association between the caffeine metabolism pathway and the phenotype of whole brain volume, diverse kernel functions yield disparate p-values, introducing ambiguity in decisions concerning null hypothesis rejection. In contrast, BayesKAT offers a more interpretable metric. (B) Based on a synthetic data cohort simulated assuming a true quadratic function, different kernel functions lead to inconsistent results. Across 500 simulation replicates, each individual kernel, including the quadratic kernel, yields inconsistent and ambiguous conclusions (barplot). In comparison, BayesKAT generates both interpretable and highly consistent results, with substantially boosted power. (C) Workflow of BayesKAT implementation for diverse types of genetic association tests to derive biological meaningful interpretations. (D) Model structures and the inference algorithms for the two BayesKAT strategies: BayesKAT-MCMC (left) and BayesKAT-MAP (right). BayesKAT-MCMC samples from posterior parameter distributions, providing a comprehensive view of the posterior parameter distributions. On the other hand, BayesKAT-MAP provides a more scalable solution, particularly well-suited for high-dimensional data.

In this study, we developed a novel **Bayes**ian **K**ernel-Based **A**ssociation **T**esting algorithm, BayesKAT (https://github.com/wangjr03/BayesKAT), which can address the kernel selection problem by selecting the optimal kernel in a data-adaptive way, while testing for the joint statistical associations of specific SNP groups with a complex phenotype. Moreover, compared to existing KBT-based methods, BayesKAT simultaneously achieves four goals in genome-wide association tests: (i) superior statistical power, by selecting the optimal kernel function based on the dataset under study; (ii) consistent results, by avoiding repeated tests based on a variety of different kernels; (iii) controlled type-1 error, without relying on unverified distributional assumptions or minimum p-value kernels; and (iv) strong computational scalability for high-dimensional and large-sample genome-wide data. Two alternative computational strategies, i.e., MCMC and MAP, are incorporated in BayesKAT, leading to additional implementation flexibilities for users. Extensively tested on a series of simulated datasets under different parameter settings, BayesKAT consistently demonstrates superior performance against existing methods. Furthermore, applied on the ADNI genotype datasets of the complex trait of whole brain volume (https://adni.loni.usc.edu), BayesKAT successfully discovered mechanistically related genes and biological pathways with higher accuracy. Specific genes and pathways related with neurodegenerative diseases, as reported by previous studies, are consistently prioritized by BayesKAT while not prioritized by other methods. Strikingly, BayesKAT is able to identify group-level SNP effects of novel co-expressed gene modules and protein complexes that potentially participate in the molecular processes modulating the whole brain volume phenotype. These algorithmic advantages and new biological discoveries robustly support the statistical innovation of BayesKAT and strongly highlight its effectiveness in decoding the genetic basis and associated molecular mechanisms underlying complex diseases.

## MATERIALS AND METHODS

### Overview of kernel based testing models for genetic data

Under the kernel machine regression framework, continuous quantitative traits can be associated to genetic variants or molecular features, along with additional covariates, through a semiparametric model:

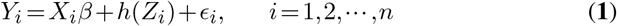

where *Y*_*i*_ denotes the continuous value of the trait for the *i*th person in a sample of size *n*; *X*_*i*_ =[*X*_*i*1_,*X*_*i*2_,*00B700B700B7,X*_*ik*_] is a set of *k* covariates for the *i*th individual that need to be controlled; and *β* =[*β*_1_,*β*_2_,*00B700B700B7,β*_*k*_] are the corresponding effects of covariates. *Z*_*i*_ =[*Z*_*i*1_,*Z*_*i*2_,*00B700B700B7,Z*_*ip*_] is the vector for the *p* genetic variants or molecular features, where *Z*_*ij*_ denotes the *j*th genetic variant or molecular level feature for the *i*th individual . The unknown errors *ϵ*_*i*_ are assumed to be independent and follow *N* (0,*σ*^2^), where the value *σ*^2^ is also unknown. The most common genetic features are SNPs and the widely used molecular-level features include gene expressions. The features, i.e. *Z*_.*j*_,*j* = 1,2,,*p* are associated with the trait, i.e. *y*, through an arbitrary function *h*(*00B7*) which is assumed to lie in a function space *H*_*K*_ generated by a kernel function *K*(*00B7,00B7*).

A kernel function is defined as a function *K* : *X ×X →* R, where the kernel matrix 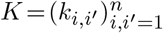 is symmetric and positive semidefinite with *k*_*i,i′*_ = *k*(*Z*_*i*_,*Z*_*i′*_). In this setting, *k*(*Z*_*i*_,*Z*_*i′*_) is a measure of similarity between the *i*th and the *i*^*′*^th subject. There are a variety of kernel functions to choose from, and the most widely used ones include the Linear kernel, the Quadratic kernel and the Gaussian kernel. For genetic SNP data, identity by state (IBS) kernel is a popular candidate kernel function suggested by various studies (19), (10), (12). The functional forms of these kernels are summarized below:

- Linear kernel: 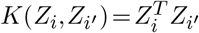
- Quadratic kernel: 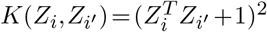
- Gaussian kernel: 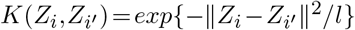 where 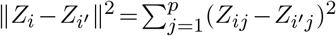 is a tuning parameter.
- IBS kernel: 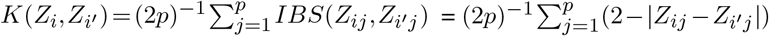

It has been shown (11) that the kernel machine regression model in equation (**1**) is equivalent to the following linear mixed model:

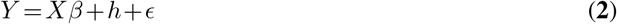

where *β∈*ℝ^*k*^ is a vector of effect sizes for covariates *X∈*ℝ^*n×k*^, *h* is an *n×*1 vector of random effects which is distributed as *h∼N* (0,*τ K*), where *K* is the *n×n* kernel matrix and the error is distributed as *ϵ∼N* (0,*σ*^2^*I*), where *σ*^2^ is the error variance and *τ* is a variance component for the genetic effect.

The main goal is to test if the genetic variants have any combined effect on the outcome variable *Y* . Testing for the presence of group effect of *Z* is equivalent to testing the hypothesis *H*_0_ : *τ* = 0 vs. *H*_1_ : *τ >* 0. Most KBT methods, including SKAT, choose the kernel function first and then perform the testing based on the specified kernel function.

### Importance of choosing appropriate Kernel

Although a variety of different kernels are available, for a given dataset, it is practically impossible to know a priori which kernel will fit the dataset best and maximize the testing power. Genetic data related to complex phenotypes pose particular challenges, primarily stemming from our limited understanding of how the interplay among genetic or molecular features influences their collective association with a phenotype. Therefore, choosing a kernel randomly can lead to a less powerful testing procedure for genome-wide applications. For example, if the outcome variable *Y* is related to the features through a Quadratic function, using a Linear kernel in the model will lead to weak tests that are not able to reject the null hypothesis even when the association is strong. On the other hand, by repeatedly applying KBT models based on different candidate kernels and choosing the one resulting in the minimum p-value, there is a high chance of making a false discovery, i.e., rejecting the null hypothesis when there is no association.

Combining a panel of candidate kernels together to create a composite kernel is thus a natural and effective strategy to overcome this issue. While a straightforward strategy of averaging kernels to form a composite kernel, i.e., a linear combination of candidate kernels with equal weights, can perform better than the worst-performing kernel function, it usually cannot perform as efficiently as the best kernel to accurately represent the association between the trait and features for a given dataset, thus, is not guaranteed to increase the statistical power. As shown in 1(A)(B), evaluated on both real and synthetic datasets, the average kernel strategy can often lead to incorrect and inconsistent results in practice. Thus, a systematic data-adaptive approach of optimal kernel selection is highly desirable for high-dimensional genome-wide association tests, especially for complex human disease phenotypes that are genetically modulated by multiple inter-dependent genetic variants.

### BayesKAT

Our new algorithm, BayesKAT (https://github.com/wangjr03/BayesKAT) employs a novel Bayesian modeling strategy to automatically select the optimal composite kernel based on the data and does not require the composite kernel function to be set a priori by the user. Based on the inferred optimal composite kernel function, BayesKAT can efficiently test the joint effects induced by a group of genetic or molecular features associated with a phenotype. The optimal composite kernel is a linear combination of candidate kernels where the weight of each candidate kernel reflects the degree of usefulness of the kernel explaining the complex relationship between a group of features and the phenotype of interest. As an illustration, suppose there are three potential kernels: Quadratic, Gaussian, and IBS. If the IBS kernel effectively captures the underlying relationship, it will carry greater significance within the composite kernel, hence have a larger weight, while the impact of other kernels may be relatively weak, as indicated by their lower weights. Figure 1(C) provides an overview of the workflow of BayesKAT and its two computational strategies: 1) the Markov Chain Monte Carlo (MCMC) strategy; and 2) the Maximum a Posteriori (MAP) strategy. Additionally, it is noteworthy that while BayesKAT is primarily developed to test genetic associations, it can also be employed for a wide range applications, including testing the association between continuous gene expression features and complex traits.

Consider a set of *m* candidate kernelLs *K*_1_,*K*_2_,*00B700B700B7,K*_*m*_, the composite kernel is in the form of 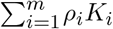 where 0 *≤ρ ≤* 1 and 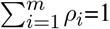 and

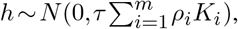

Therefore, selecting the optimal composite kernel is equivalent to selecting the optimal value for the weight *ρ*_*i*_ (*i* = 1, *00B700B700B7,m*) so that it can capture the underlying relationship between the genetic or molecular features and the trait, when testing the group-level effect of a set of multiple features.

#### Input data organization and model set up for BayesKAT

Data containing the information of genotypes or molecular features for complex phenotypes can be collected from large public-accessible or user-generated cohorts (e.g., ADNI, UK biobank, All of Us (https://allofus.nih.gov/), GTEx, PsychENCODE (https://psychencode.synapse.org/), etc.). Individual-level data can be pre-processed and efficiently undergo steps of quality controls using software such as Plink (26). Additionally, biological meaningful feature groups need to be defined and created depending on the goals of genome-wide association tests. In this study, we have explored four different biology inspired ways of grouping functionally related genetic variants, including 1) gene-wise groups: aggregating SNPs located within genomic regions of genes; 2) pathway-level groups: aggregating SNPs situated within genes that belong to a specific molecular pathway; 3) co-expression gene modules: aggregating SNPs located within genes that belong to a specific co-expression module; and 4) Protein-protein interaction(PPI) modules: aggregating SNPs located within genes that belong to a specific PPI module, which may represent a protein complex.

As a kernel-based testing model, BayesKAT relies on a set of candidate kernel functions, which are incorporated to infer the optimal composite kernel for the association tests. For the convenience of practical implementations, BayesKAT infers the optimal composite kernel consisting of three candidate kernels as the default setting. And the default candidate kernels include Quadratic, Gaussian and IBS kernel. To construct a composite kernel, the candidate kernels are normalized in BayesKAT based on the previously proposed technique (21) so that they are in the same scale and comparable.

To gain robust performance, weakly informative prior distributions are used for model parameters by default, although the users can incorporate more informative priors based on specific knowledge about the data. The important model parameters are 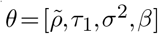, where 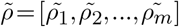 are the unscaled weights of the candidate kernels 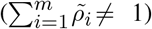, *β* =(*β*_1_,*β*_2_,*00B700B700B7,β*_*k*_)^*T*^ and 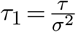 after reparameterization. And the weakly informative prior distributions are:

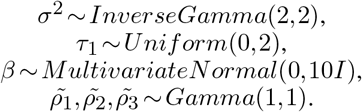

The actual weights for candidate kernels 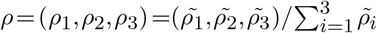 Clearly 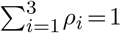 and *ρ∼ dirichlet*(1,1,1)

The data distribution *y*|*θ∼N* (*Xβ,σ*^2^(*τ*_1_*K*_*c*_ +*I*)), where the composite kernel 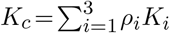

#### BayesKAT strategy

Here the main hypothesis to test is: *H*_0_ : *τ* = 0 vs. *H*_1_ : *τ >* 0. It is equivalent to test *H*_0_ : *τ*_1_ = 0 vs. *H*_1_ : *τ*_1_ *>* 0. Bayes factor(*BF*_10_) is calculated to test the hypothesis, which evaluates the evidence in favor of the alternative hypothesis. Bayes factor is defined as the ratio of marginal likelihoods under two hypotheses (27):

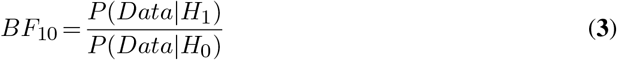

where *P* (*Data H*_0_) and *P* (*Data*|*H*_1_) are the marginal likelihoods under *H*_0_ and *H*_1_, respectively. Given the input data, BayesKAT mainly uses two efficient and easy-to-implement strategies to select the composite kernel and calculate the Bayes Factor *BF*_10_ by estimating *P* (*Data*|*H*_0_) and *P* (*Data*|*H*_1_). The two computational strategies are explained in subsequent sections.

#### Interpreting BayesKAT output

The preliminary output from BayesKAT, *BF*_10_, is a summary of evidence provided by the data in favor of *H*_1_ as opposed to *H*_0_. In addition, the posterior probability of the association (*P* (*H*_1_|*Data*)) is calculated as the final output, which has a one-one relation with *BF*_10_, i.e.,

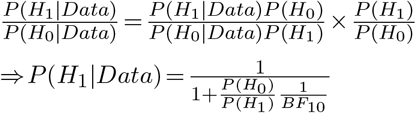

Here *P* (*H*_1_) and *P* (*H*_0_) are the prior probabilities under *H*_1_ and *H*_0_, respectively, i.e. the probabilities of existence of association and no association, respectively. Depending on the specific biological problem and dataset, the values of *P* (*H*_0_) and *P* (*H*_1_) can be set given biological evidence or prior knowledge, and *P* (*H*_1_|*Data*) can be calculated. *P* (*H*_1_|*Data*) is the posterior probability of the model under *H*_1_ given the data. *P* (*H*_1_|*Data*) gives a quantitative evaluation of how probable there exists an association between the genetic features and the phenotype of interest or how strong the evidence is against the null hypothesis.

As already demonstrated by the example in Figure 1(A,B) in the introduction section, existing methods based on pre-selected kernels or average composite kernels suffer from inconsistent hypothesis testing results and may lead to false discoveries. In contrast, for the Figure 1A scenario, BayesKAT calculated the posterior probability of a true association given the data P(*H*_1_|*data*)=0.19, i.e. the evidence of association is very low and the existence of true association is not very likely. This is consistent with the fact that there is a lack of documented evidence of the association between the caffeine metabolism pathway and the whole brain volume phenotype. Strikingly, in the scenario of the synthetic data cohort simulated with known association based on the Quadratic kernel (see Figure 1(B)), BayesKAT successfully calculated the posterior probability *P* (*H*_1_|*Data*) = 0.99 to suggest the existence of association, without incorporating any prior information. Moreover, tested on 500 repeated simulation cohorts, BayesKAT achieves much higher statistical power than other methods and also demonstrates more consistent testing results (see Figure 1(B)). Investigating the inferred weights for each candidate kernel functions in the final composite kernel selected by BayesKAT further shows that the Quadratic kernel is correctly assigned with the largest weights (Supplementary Figure 1), suggesting that BayesKAT can efficiently capture the functional form of the underlying statistical associations in a data adaptive way.

### BayesKAT-MCMC

As a Bayesian model, BayesKAT-MCMC employs the Markov Chain Monte Carlo (MCMC) sampling-based strategy to infer the optimal composite kernel function. Leveraging MCMC for efficient and traceable samplings from complex target distributions, BayesKAT-MCMC avoids direct sampling from the posterior distribution *P* (*θ*|*Data*) (see section “*Input data organization and model set up for BayesKAT*”), which does not have a closed mathematical form and is computationally intractable. Instead, Metropolis-Hastings method (28, 29) is used to iteratively draw samples based on the generated Markov chain, which are able to approximate the target probability distribution *P* (*θ*|*Data*).

Let *θ*_0_ denote the initial value for *θ*. The *t*th iteration of the Metropolis-Hastings algorithm consists of the following steps (30), (31):

1. Sample a candidate point *θ*_*t*_from a proposal distribution *J*_*t*_(*θ*_*∗*_|*θ*_*t−*1_).
2. Calculate the acceptance ratio for jumping to the new point 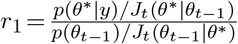
3. Set *θ*_*t*_ = *θ*^*∗*^ with probability *r*_1_ and *θ*_*t*_ = *θ*_*t-*1_ with probability 1-*r*_1_. That is, it jumps to the new proposed value with probability *r*_1_ and stays at the same value with probability 1*−r*_1_.

BayesKAT-MCMC uses the R package BayesianTools (32) to generate two sets of samples from the posterior distribution of *θ* under the hypotheses *H*_1_ and *H*_0_, using the Metropolis-Hastings algorithm in an adaptive way (33) to leverage the history of the stochastic process and appropriately fine-tune the proposal distributions. Three separate MCMC chains initiated from different random start points are generated for 50,000 iterations. To ensure that the MCMC chains are converged, trace plot is used to visualize the moves of the Markov chains in the state space (34) . In addition, based on the Gelman-Rubin diagnostic method (35), the potential scale reduction factor, i.e. PSRF score, is also calculated and presented at the end of MCMC sampling to inspect whether the chain is converged in which the PSRF score is close to one.

Based on the generated posterior samples, the marginal distributions of the parameters are further visualized as shown in Figure 1(D). The posterior samples are used to estimate the composite kernel weights and also the marginal likelihoods *P* (*Data*|*H*_*i*_),*i* = 1,0 using the Chib’s method (36). Bayes Factor is subsequently calculated using the formula presented in equation (**3**).

### BayesKAT-MAP

Although BayesKAT-MCMC yields comprehensive information of the marginal distributions of model parameters, drawing large sets of samples from the posterior distributions in the MCMC strategy is computationally expensive. Here, we provide an alternative strategy, termed BayesKAT-MAP, which is easy to implement, allows parallel calculations,and has higher computational scalability. Instead of drawing numerous MCMC samples from the posterior distributions and then estimating the parameter values, BayesKAT-MAP employs the quick optimization technique to estimate the parameters of interest directly, based on the Maximum A Posteriori (MAP) strategy such that,

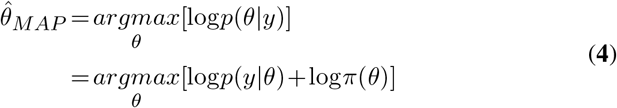

where *π*(*θ*) denotes the prior distribution *θ*. Because the objective function is nondifferentiable at some points, a derivative-free optimization algorithm by Quadratic approximation using the R package Minqa (37) is implemented. The most important model parameter *τ*_1_ indicates the existence of association between the feature set and the trait variable, with *τ*_1_ = 0 suggesting that there is no association. In BayesKAT-MAP, if the calculated MAP estimator of 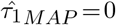, it follows that the Bayes Factor =0 (i.e., no evidence of association is found), and the computational process terminates. On the other hand, if 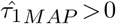, it implies that there might be some evidence of association and BayesKAT-MAP proceeds to calculate the MAP estimator again under *H*_0_ and then computes the marginal likelihoods *P* (*Data*|*Hi*),(*i* = 1,0), along with the Bayes Factor.

Due to the practical limitations of exact analytical methods, such as relying on specific distributional assumptions, efficient numerical integration approaches (38) are needed to calculate the marginal likelihoods under hypotheses *H*_*i*_, *P* (*Data*|*H*_*i*_) = *Pr*(*Data*|*θ,H*_*i*_)*π*(*θ*|*H*_*i*_)*dθ*, so that the model can be applied on diverse panels of data. BayesKAT-MAP employs the Laplace’s method (39), (40), (27) for approximating the integral *T* = *Pr*(*Data*|*θ,H*_*i*_)*π*(*θ*|*H*_*i*_)*dθ* by 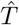, where

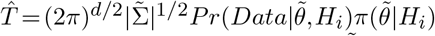

where *d* is the dimension of *θ*, 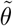 is the mode of the log-likelihood function *l*(*θ*|*Data*), 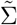 is the inverse of the negative Hessian matrix of the second derivative of *l*(*θ*|*Data*) computed at 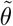. For boundary regions of the parameter space, the Laplace approximation is modified according to the previously developed protocol (41) to accommodate the boundary cases. Based on the estimated marginal likelihood densities, the Bayes Factor is then computed and the posterior probabilities are finally inferred, given user-defined priors *p*(*H*_0_) and *p*(*H*_1_) for which non-informative priors are used as the default setting in BayesKAT.

### Comaparison between BayesKAT-MCMC and BayesKAT-MAP

The performance and runtime of BayesKAT-MCMC and BayesKAT-MAP under different settings are systematically compared. Table 1 summarizes the empirical type-1 error and empirical power for different simulated functional dependencies. The simulations were based on the proposed settings used by previous studies (21). As shown in Table 1, both MCMC and MAP strategies achieve nearly equal statistical power in detecting associations. While BayesKAT-MCMC provides more information on the posterior distributions of parameters, it is more computationally expensive compared to BayesKAT-MAP. When the number of samples or features, i.e. *n* or *p*, increases beyond 500, BayesKAT-MCMC is not sufficiently scalable without requesting more high-performance computing resources. On the other hand, Figure 1(D) shows the superior computational scalability of BayesKAT-MAP based on the same level of computational resource support. Therefore, these two alternative strategies provide similar accuracy and complementary signatures for genetic association tests of complex traits, with BayesKAT-MAP being more flexible and conservative. For the rest of the paper, results of BayesKAT-MAP are presented due to its scalability. The code for implementing BayesKAT-MCMC and BayesKAT-MAP are both made publicly available via GitHub: https://github.com/wangjr03/BayesKAT.

**Table 1.** Empirical power of BayesKAT-MCMC and BayesKAT-MAP.

| $h(Z)$ | $n$ | $p$ | BayesKAT-MCMC | BayesKAT-MAP |
| --- | --- | --- | --- | --- |
| $h(Z)=0$ | 500 | 500 | 0.018 | 0 |
| $h(Z)=2 \times Z_1 Z_3$ | 500 | 500 | 0.946 | 0.946 |
| $h(Z)=2 \times Z_1 Z_3 + 0.04 \times Z_i + 0.04 \times Z_3$ | 500 | 500 | 0.948 | 0.958 |
| $h(Z)=0.4 \times (Z_1 - Z_3) + 0.4 \times \cos(Z_3) \exp(-Z_3^2/5)$ | 500 | 500 | 0.976 | 0.998 |

The functional forms in simulation scenarios are taken from (21) and coefficients are adjusted based on $n$ (sample size) and $p$ (no. of features).

### Comparison with other methods and performance evaluation

The performance of BayesKAT is compared to two state-of-the-art algorithms: 1) SKAT using the average composite kernel, denoted as SKAT(Avg) (18); and 2) the U statistic-based method, denoted as mKU (21). Both of these two methods are frequentist approaches that maximize the power after restricting the type 1 error to a fixed level of *α*, such as setting *α* to 0.05, and have been shown to outperform other existing methods. In contrast, as the first Bayesian model for this problem, BayesKAT uses a fixed threshold on the posterior probability or the Bayes factor, based on previously suggested guidelines (27), to reject the null hypothesis. To make fair comparisons, the performance of each method (i.e., the empirical statistical power), is evaluated at a fixed and equal empirical type 1 error across all three algorithms. A systematic comparison based on rigorous simulations are presented in the Results section.

### Multiple testing correction

When *m*_1_ multiple groups are simultaneously tested, multiplicity corrections are needed. Multiple testing corrections on p-values from frequentist models are carried out using methods such as the Bonferroni correction (42), which controls the family-wise type 1 error to *α* by setting the individual test’s type 1 error at *α/m*_1_. Other methods like FDR control have also been popular ones. For multiple testing corrections on Bayesian models, the multiplicity control is achieved by setting a high prior probability of the individual null hypothesis, as suggested by previous studies (43). For each test, the prior probability is set as *P* (*H*_0_) = 0.99, which is equivalent to assuming that, on average, one in 100 tests is believed to have a true association. Considering the number of SNP groups in genetic applications, such as the number of biological pathways or co-expression modules, this choice of prior probability setting is regarded as rather conservative and ensures fair performance comparisons.

### Real data preprocessing

In addition to a series of simulated datasets, real genetic datasets are used to evaluate the performance of BayesKAT and its derived biological discoveries. The data used in the preparation of this article were obtained from the Alzheimer’s Disease Neuroimaging Initiative (ADNI) database (adni.loni.usc.edu). The ADNI was launched in 2003 as a public-private partnership led by Principal Investigator Michael W. Weiner, MD. The primary goal of ADNI has been to test whether serial magnetic resonance imaging (MRI), positron emission tomography (PET), other biological markers and clinical and neuropsychological assessments can be combined to measure the progression of mild cognitive impairment (MCI) and early Alzheimer’s disease (AD). Plink software (26) (http://pngu.mgh.harvard.edu/purcell/plink/) is used to pre-process the individual-level genotype data.

Four different biology-based strategies are used to create the feature groups of functionally related SNPs for genome-wide testing. The group-level statistical test results based on these different SNP grouping strategies lead to complementary biological insights into the genetic basis underlying the specific phenotype. The four SNP feature grouping strategies are: (1) Gene-wise SNP groups for 18,999 protein-coding genes in the human genome(44). For each protein-coding gene, all SNPs located within +/-5KB of the gene body are collected as the gene-wise SNP feature set; (2) Pathway-wise SNP groups for 352 KEGG pathways (23), (24), (25). For each pathway, gene-wise SNP groups for all genes belonging to the specific pathway are collected as the pathway-wise SNP feature set. The number of SNPs per pathway varies from 35 to 22,555, with a mean of 1,721 SNPs. Figure 5(A) provides a schematic figure demonstrating the pathway-wise joint SNP testing procedure; (3) Co-expression gene module based SNP groups. Forty-one co-expression gene modules are identified using the R package “WGCNA” (45), (46) to find correlated gene co-expression clusters from expression data (available on the ADNI website). The co-expression gene modules have a different number of genes in them, which varies from 6 to 3361. SNPs within each module are extracted for further group-wise testing; and (4) 401 protein-protein interaction (PPI) gene module based SNP groups. The PPI gene modules are previously created in (47) based on the topology of the PPI network. The number of genes in PPI modules ranges from 2 to 497.

## RESULTS

### Enhanced efficacy of BayesKAT benchmarked on simulation studies

As defined in the section of Materials and Methods, the rows of the feature matrix *Z∈*ℝ^*n×p*^ correspond to *n* individuals and the columns correspond to the *p* features. Depending on the particular genetic association studies, the features may encompass discrete genetic characteristics, like alleles with values of 0, 1, or 2, or continuous molecular features, such as gene expressions. We have conducted simulations for both cases, with *Z* corresponding to discrete or continuous features, under both low and high dimensional settings.

#### Simulation with continuous features

As shown in Figure 2(A), the simulations based on continuous features are first conducted to evaluate the performance of BayesKAT using similar scenarios presented in (21). With the specified parameters (*n* = 500, *p* = 100, *k* = 2, *m* = 3), the feature matrix *Z* is simulated from a multivariate normal distribution with mean 0_*p*_ and an AR(1) correlation matrix *R* where (*R*(*j,j*^*′*^) = *r*|*j−j*_*′*_|). In the simulation, the correlation *r* is set to be 0.6. The covariate matrix *X∈* ℝ_*n×*2_ has one binary covariate generated from a Bernoulli (0.6) distribution and one continuous covariate generated from N(2,1). The covariate coefficients are set as *β* =(0.03,0.5) and the outcomes *Y* are simulated based on model (**1**). The error term with variance *σ*^2^ = 1 is added as the random noise. Three commonly used candidate kernels are incorporated in BayesKAT: Linear, Quadratic, and Gaussian. Different functional forms of *h*(·) are used to create different scenarios to test the performance. The empirical type 1 errors of each method are calculated based on the specific scenario where *h*(*Z*) = 0, i.e. there is no association between *Z* and *Y* in the simulated data. The different scenarios are given below:

- Scenario A: *h*(*Z*) = 0.6*×Z*_1_*Z*_3_
- Scenario B: *h*(*Z*) = 0.55*×Z*_1_*Z*_3_+0.1*×Z*_1_ +0.1*×Z*_3_
- Scenario 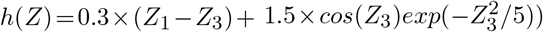

In all the simulation scenarios, as shown in Figure 2(B), the empirical power versus empirical type 1 error for BayesKAT is consistently better than that of SKAT(Avg) and mkU. Moreover, sensitivity analyses are conducted to evaluate the effects with different simulation parameters on the final performance of different algorithms. Notably, when the number of features *p* increases from 100 to 150, while keeping all other parameters fixed, BayesKAT consistently achieves superior empirical power compared to other methods (see Figure 2 (B)). Similarly, when the correlation *r* increases from 0.6 to 0.8 with *p* fixed as 150, BayesKAT consistently demonstrates higher empirical power and outperforms other methods. Taken together, these simulation results demonstrate the robust superior performance of BayesKAT compared to SKAT(Avg) and mKU in various settings with continuous features.

**Figure 2.**
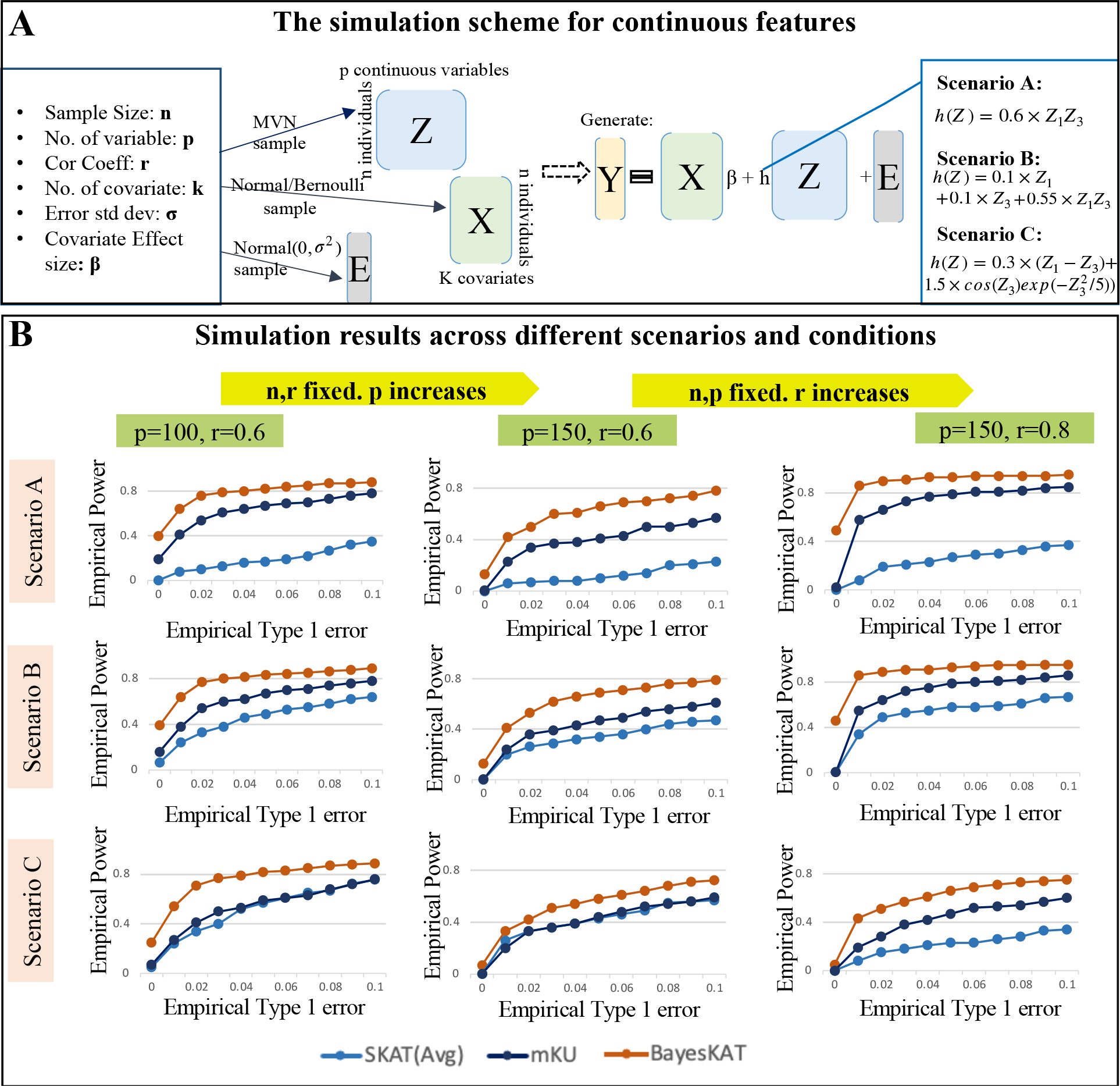
Performance comparison based on simulations using continuous features. (A) Schematic summary of the data generation process for continuous molecular features (e.g. gene expression features) and the demonstration of the implementation under various scenarios. (B) Performance comparison across different simulation settings with systematic performance evaluations, i.e. the empirical power versus empirical type 1 error, for SKAT(Avg), mKU, and BayesKAT across different scenarios and parameter settings. When increasing *p* while keeping other factors constant, all methods exhibit a slight decline in power, but BayesKAT consistently outperforms SKAT(Avg) and mKU. Additionally, as *r* increases under fixed parameters, BayesKAT also consistently surpasses SKAT(Avg) and mKU.

#### Simulation with discrete features

The performance of different models on discrete SNP features is first evaluated based on simulations where randomly selected SNP groups are used as features. Because randomly selected SNPs are generally not functionally related, the overall effectiveness of all KBT models decreases as expected, with BayesKAT still showing improved empirical power compared to other methods (Supplementary Figure 2). Because the real-world implementations of KBT models for genetics studies usually focus on functionally related groups of SNPs, a more realistic strategy of simulating groups of discrete SNP features, instead of randomly selected unrelated SNPs, is employed to further benchmark the performance of BayesKAT (Figure 3(A)).

**Figure 3.**
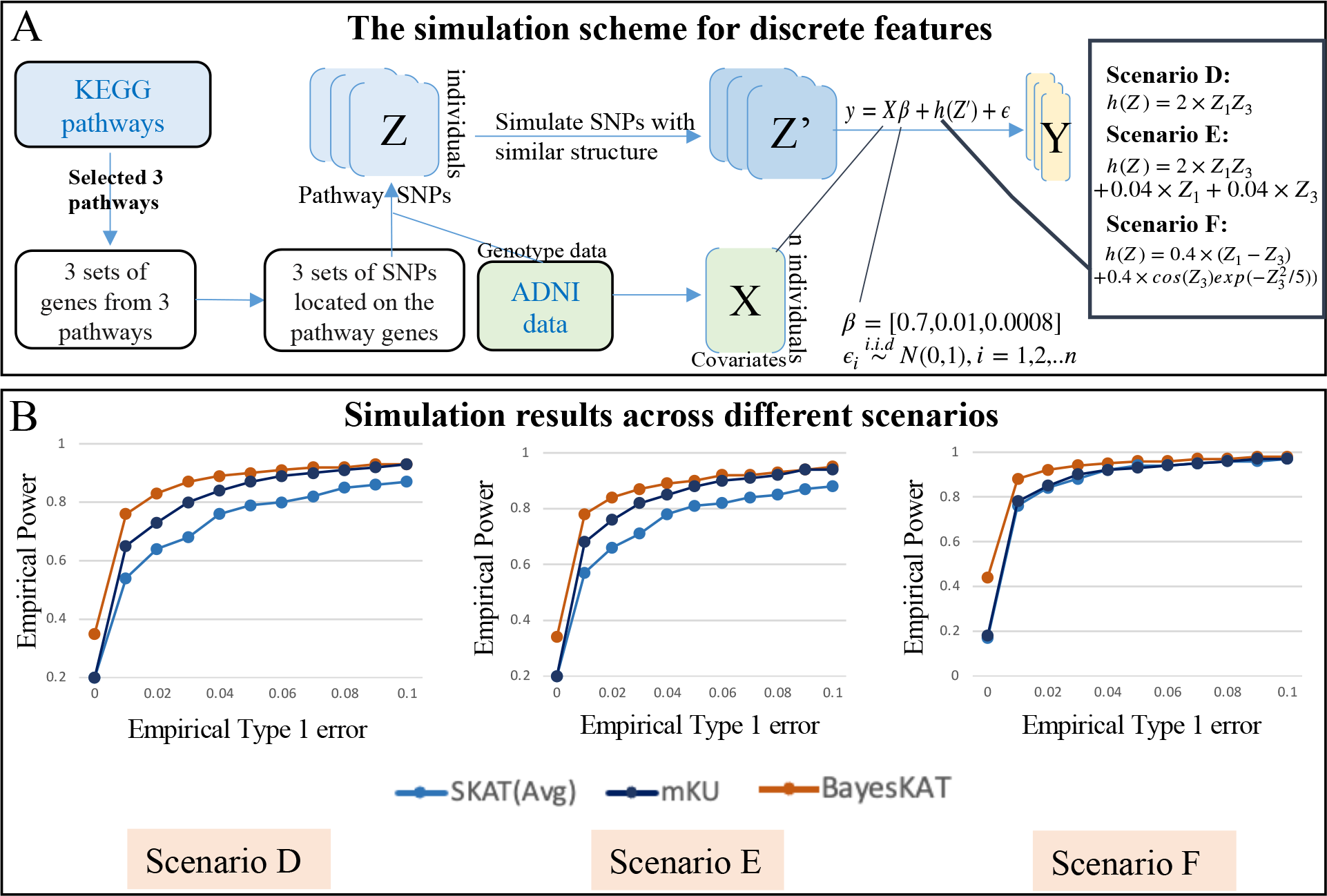
Performance comparison based on simulations with discrete features. (A) Schematic summary of the data generation process for discrete genetic features (e.g., SNP variants) and the demonstration of the implementation under various scenarios. Starting with the original *Z* matrix containing groups of SNPs from each pathway, a simulated SNP matrix *Z*^*′*^ is generated, preserving the underlying interrelationships among the SNPs. Covariate variables (i.e., age,gender and occupation) are incorporated based on the data from ADNI. (B) Performance comparison across different simulation settings. Systematic performance evaluations, i.e. the curves of empirical power versus empirical type 1 error, for SKAT(Avg), mKU, and BayesKAT across different scenarios are plotted. The performance across three sets of simulated datasets are averaged. The performance curves based on each individual simulated dataset can be found in Supplementary Figure 3.

To rigorously capture the underlying linkage disequilibrium structures of discrete features in real-world SNP data, the ADNI dataset is used as the basis for the simulations, where the covariate matrix *X* is created from the real covariates (e.g. age, gender and occupation) of the corresponding individuals in the dataset, and SNPs located in specific genes belonging to the selected KEGG pathways are included into the testing (see Materials and Methods). Three different KEGG pathways are randomly selected for performance evaluations. For each pathway, the groups of SNPs located in the corresponding gene members are identified. The “SNPknock” package (48) is then used to simulate the knockoff SNP data, which maintains the structural dependency among SNPs in each group intact in the simulated knockoff SNPs (Figure 3(A)). The feature matrix *Z* is constructed based on the simulated SNP data, with *n* = 755 and *p* ranging between 4000 and 5000. The outcome variable *Y* is simulated based on three different scenarios. Each scenario corresponds to a different functional form *h*(*Z*), that is:

- Scenario D: *h*(*Z*) = 2*×Z*_1_*Z*_3_
- Scenario E: *h*(*Z*) = 2*×Z*_1_*Z*_3_ +0.04*×Z*_*i*_ +0.04*×Z*_3_
- Scenario 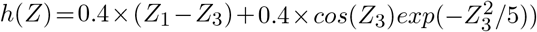

Where *Z*_*i*_ is the *i*th column of *Z* corresponding to the *i*th SNP. For each group of pathway-level knockoff SNPs, 500 simulations are generated. By applying BayesKAT and the other methods on the set of simulations, the corresponding empirical type 1 error and empirical power are calculated accordingly. The summary of performance comparisons based on this extensive set of simulations is shown in Supplementary Figure 3. The empirical power and empirical type 1 error for each SNP group clearly demonstrates that BayesKAT robustly outperforms SKAT(Avg) and mKU, across different simulation scenarios and settings. The averaged performance over 3 pathway-level SNP groups can be found in Figure 3(B). Remarkably, in addition to these consistent advantages, BayesKAT also achieves much lower empirical type 1 error across all simulation settings, when the suggested Bayes Factor threshold (27) is employed. It suggests that the associations between the SNP groups and the phenotype detected by BayesKAT exhibit a significantly higher level of reliability compared to other methods, a crucial attribute for genetic applications in complex diseases.

### BayesKAT reveals novel associated genetic basis of complex traits

To illustrate the novel biological insights that can be generated by BayesKAT, the individual-level data, including the genotype, phenotype and demographic covariates, from the ADNI project (https://adni.loni.usc.edu) (49), (22) are used to conduct a series of group-level genetic association testings. Specifically, BayesKAT is used to test the group-wise associations between SNP sets and the complex phenotype of whole brain volume, based on the available information across 755 individuals in the ADNI cohort.

#### BayesKAT prioritizes functionally related genes

To identify genes associated with the trait of whole brain volume, we conduct a gene-wise association test with gene-level SNPs (see Materials and Methods). BayesKAT prioritized 17 genes, whose posterior probability of association (*P* (*H*_1_|*Data*)) is greater than 0.7. Figure 4(A) shows some examples of the prioritized genes, which have been suggested to be associated with brain or neurodegenerative disorders by previous seminal studies (50), (51), (52). The whole list of the 17 prioritized genes and their corresponding posterior probability of associations are provided in Supplementary Table 1. Strikingly, as external evidence in support of BayesKAT’s prioritized genes, 12 out of the 17 genes (71%) contain CpGs that have been found to be involved with significant meQTLs (53) in the human brain cortex. In contrast, the genes prioritized by SKAT(Avg) and mKU demonstrate much lower fractions of overlapping with CpGs linked to meQTLs (66% and 61% respectively, Figure 4(B)). The higher proportion of prioritized genes containing CpGs offers molecular-level support for BayesKAT’s ability to uncover genes mechanistically linked to complex traits.

**Figure 4.**
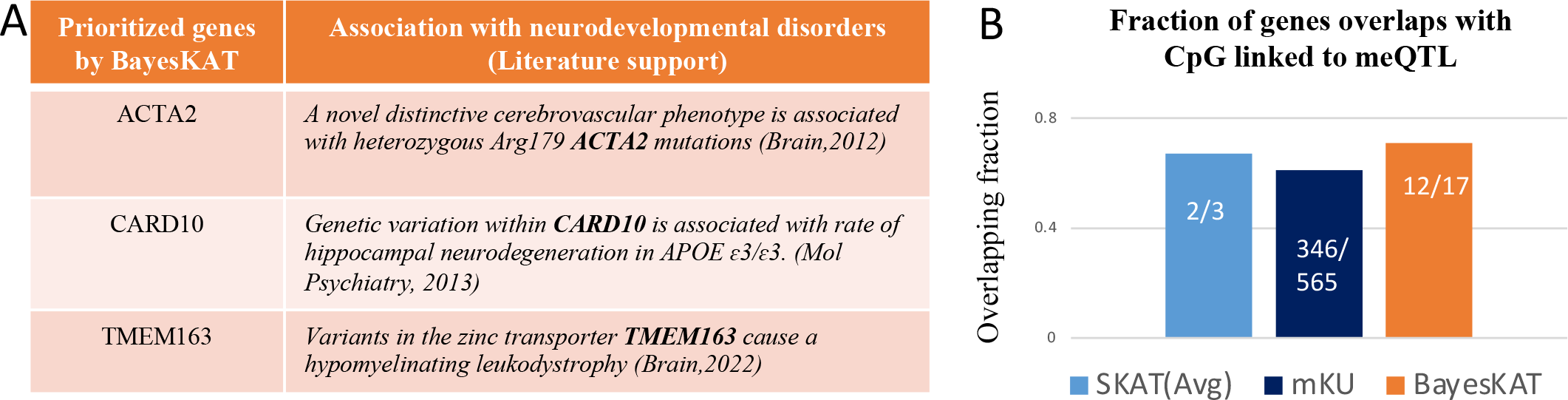
Functional validation of BayesKAT’s prioritized genes using orthogonal information. (A) Top-ranking genes prioritized by BayesKAT are strongly supported by previous literature of functional studies of brain-related diseases. (B) The selected genes by BayesKAT demonstrate higher fractions of overlapping meQTL’s CpG sites than the genes selected by SKAT(Avg) and mKU. The CpG sites of significant meQTLs from the brain tissues represent orthogonal molecular-level evidence in support of the gene’s functional involvement with whole brain volume.

**Figure 5.**
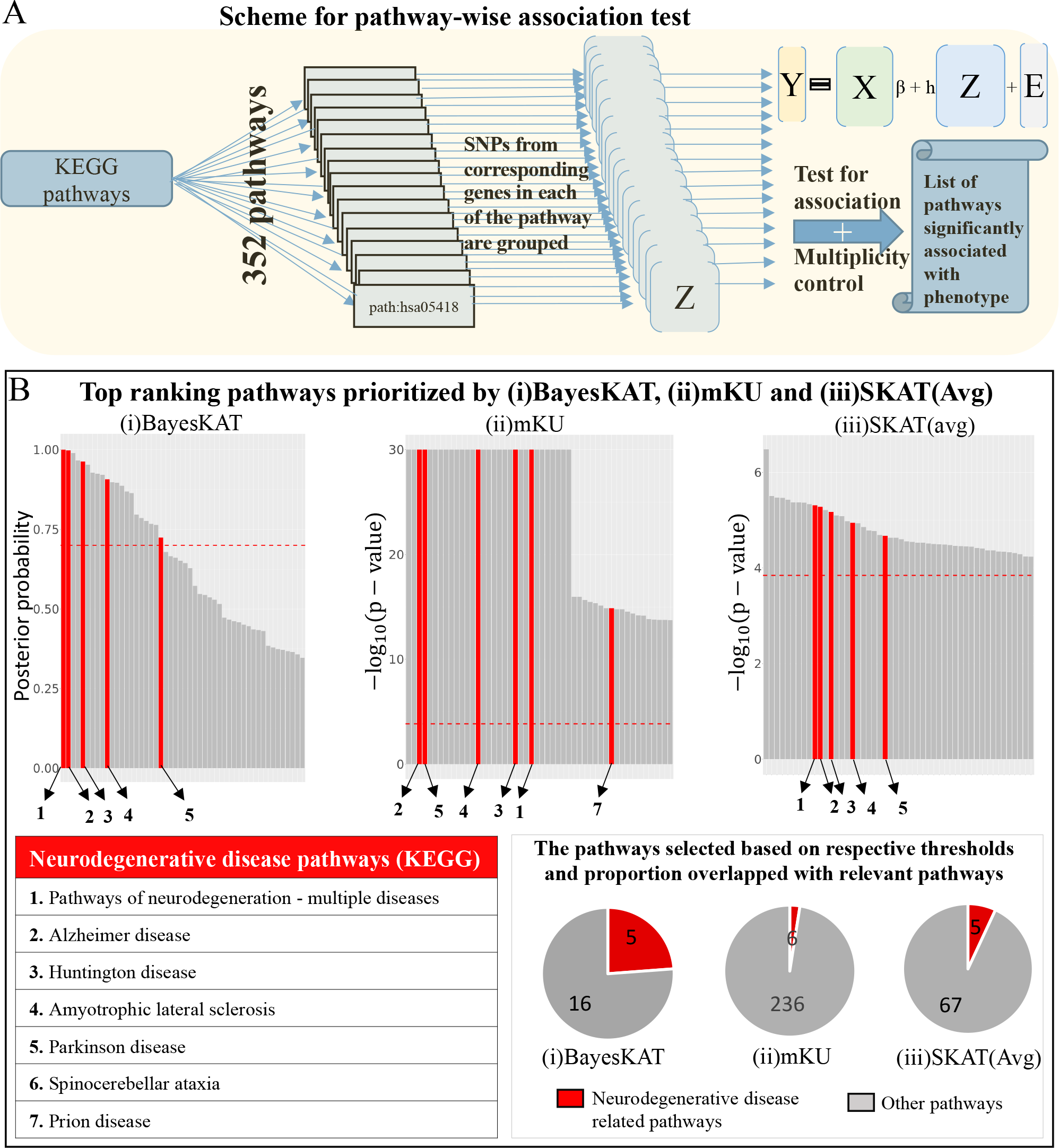
Pathway-level association tests by BayesKAT prioritizes neurodegenerative disease related pathways. (A) Schematic representation illustrating the steps of pathway-level association tests. Sets of SNPs located within genes belonging to each of the 352 pathways are tested simultaneously for pathway-level associations with the phenotype of interest (e.g., the whole brain volume). Multiplicity control is implemented to identify the specific list of significant pathways linked to the phenotype. (B) The top-ranking pathways prioritized by (i) BayesKAT, (ii) mKU, and (iii) SKAT(Avg) demonstrate distinct enrichment with neurodegenerative disease associated pathways. Top 50 pathways are shown for fair comparison. The top-ranking pathways by BayesKAT are ranked by the estimated posterior probabilities of the pathway-level associations. The top-ranking pathways by the frequentist methods, mKU and SKAT(Avg) are ranked by the -log_10_(p-values). The pathways highlighted in red are neurodegenerative disease related pathways. The red horizontal dashed line in each bar plot indicates the threshold used by each model for fair comparison (see Materials and Methods). The pie charts illustrate the proportion of the selected pathways (above model’s thresholds) that belong to the neurodegenerative disease pathways. BayesKAT notably exhibits enhanced prioritization of neurodegenerative disease pathways. Due to the issue of inflated p-values in mKU, pathways with p-values of 0 are assigned -log_10_(p-values)=30 for visualizations.

#### Biological pathways linked to neurodegenerative diseases are top-ranked by BayesKAT

To identify biological pathways that potentially modulate the whole brain volume trait, BayesKAT is used to analyze pathway-level SNP groups (see Materials and Methods, Figure 5(A)). The top 50 ranked KEGG pathways by each model are summarized in Figure 5(B), where BayesKAT ranks the pathways based on decreasing posterior probability of association (*P* (*H*_1_|*Data*)) while the mKU and SKAT(Avg) methods rank the pathways based on decreasing -log_10_(p-value). Interestingly, BayesKAT successfully prioritized most of the neurodegenerative disease related pathways with top ranks (Figure 5(B)). This is a strong mechanistic support to BayesKAT’s results, because the neurodegenerative diseases, including Alzheimer’s disease, Huntington’s disease, Amyotrophic lateral sclerosis and Parkinson’s disease, have been found to be strongly related to brain volume loss (54), (55), (56), (57). Based on a reasonable posterior probability threshold 0.7, there are 21 pathways identified by BayesKAT (Figure 5(B), Supplementary Table 2), among which five pathways are associated with neurodegenerative diseases. In comparison, the mKU and SKAT(Avg) models prioritized large numbers of pathways (242 and 72 respectively), while only a small fraction of them are neurodegenerative disease related pathways (6 and 5, respectively). For SKAT(Avg), these functionally related pathways are not even the top-ranked ones. As summarized in the corresponding pie charts in Figure 5(B), BayesKAT achieves the highest efficiency in prioritizing important pathways and resulting in fewer false discoveries than the other methods. These results are also consistent with the larger type 1 errors of mKU and SKAT(Avg) observed from simulation analyses described above. Note that, setting the posterior probability threshold is subjective, same as selecting type 1 error threshold or FDR threshold (0.05 or 0.01 or 0.1). Opting for a threshold greater than 0.7 (such as 0.9, 0.95, or 0.99) is also effective, as BayesKAT efficiently prioritizes the top pathways. Overall, the highly prioritized functional relevant pathways imply the novel biological insights that can be generated by using BayesKAT.

**Table 2.** Empirical type 1 error comparison for SNP simulation.

| Method | Error control level | Avg empirical type 1 error |
| --- | --- | --- |
| SKAT(Avg) | Error controlled at 0.05 | 0.01 |
| mKU | Error controlled at 0.05 | 0.047 |
| BayesKAT | No error controlled method | 0.002 |

To further evaluate the performance of BayesKAT in determining the optimal kernel weights, 10 randomly chosen pathways are used to test the pathway-level associations. The resulting composite kernels are compared to the results of using SKAT based on individual kernels separately. The corresponding -log_10_(p-values) metrics from SKAT using individual kernels are compared to the inferred kernel weights in the composite kernels from BayesKAT. As shown in Figure 6(A), the high similarity between the two heatmaps indicates that BayesKAT can efficiently select the optimal composite kernel automatically from the data, without relying on prior knowledge or repetitively trying different individual kernels.

**Figure 6.**
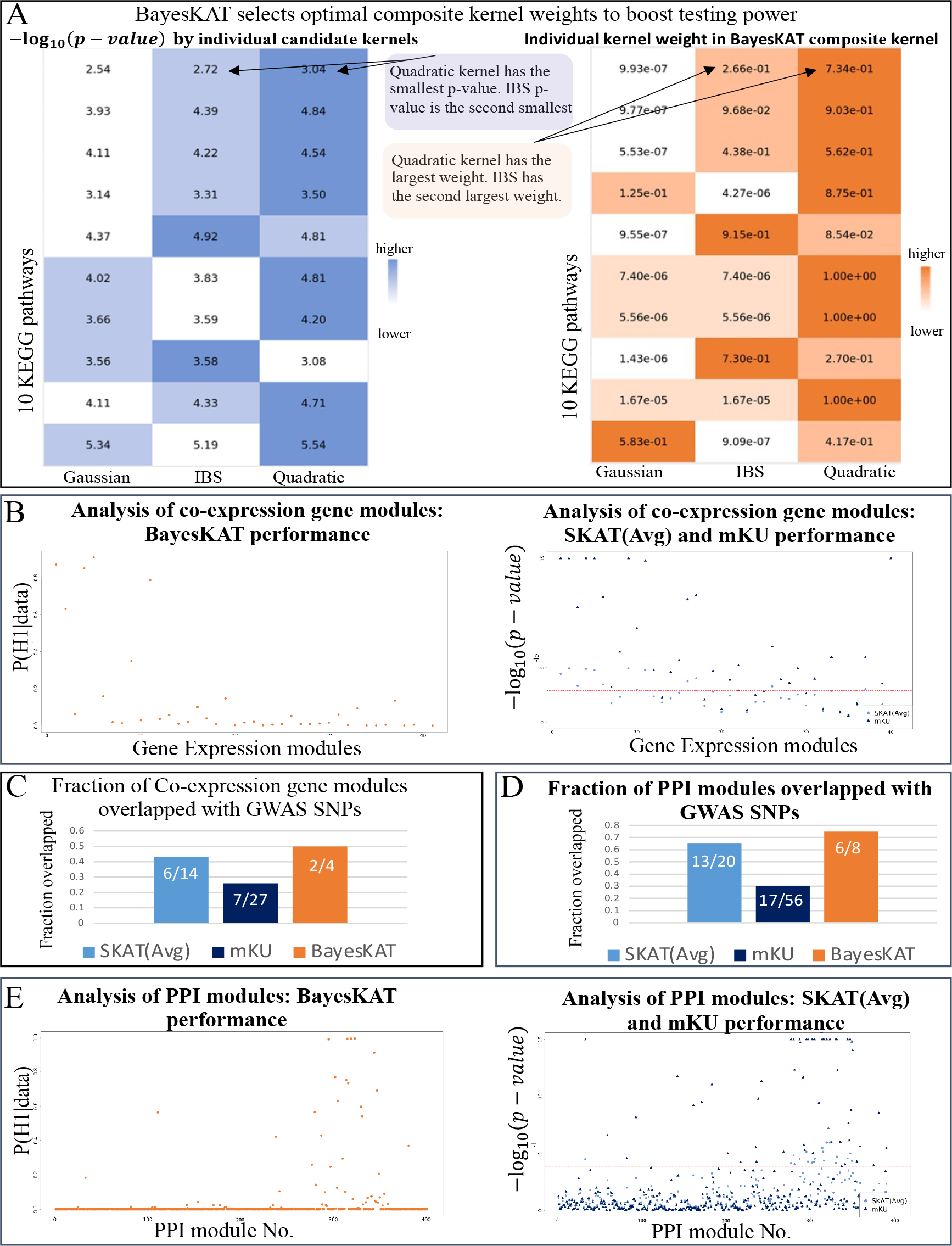
Boosted association tests based on BayesKAT’s composite kernels reveal novel modules of genes and proteins linked to brain volume. (A) The inferred weights of individual kernels in BayesKAT’s composite kernels (right) recapitulate the strength of each kernel (-log_10_(p-values)) when each kernel is incorporated separately (left). Without relying on prior knowledge or repetitively testing different kernels separately, BayesKAT automatically infers the appropriate composite kernels to boost the group-level tests for different pathways. (B) Prioritized co-expression gene modules by BayesKAT (left) vs. SKAT(Avg) and mKU (right). The significance threshold of selection for each model is represented by the horizontal red dashed lines (see Materials and Methods). (C) The selected significant co-expression gene modules by BayesKAT demonstrate higher fractions of overlapping with significant SNPs from orthogonal GWAS studies, compared to the results from SKAT(Avg) and mKU. (D) The selected significant PPI modules by BayesKAT demonstrate higher fractions of overlapping with significant SNPs from orthogonal GWAS studies, compared to the results from SKAT(Avg) and mKU. (E) Prioritized PPI modules by BayesKAT (left) vs. SKAT(Avg) and mKU (right). The significance threshold of selection for each model is represented by the horizontal red dashed lines (see Materials and Methods).

#### BayesKAT identifies trait-associated gene modules and protein complexes

To further demonstrate BayesKAT’s capability of revealing novel group-level associations to traits from specific sets of cooperative SNPs, two additional SNP grouping strategies are applied: 1) SNPs from co-expression gene modules; and 2) SNPs from protein-protein interaction (PPI) modules (see Materials and Methods). Applied on the SNP groups aggregated from co-expression gene modules, BayesKAT is able to pinpoint specific modules as significantly associated with the whole brain volume trait (Figure 6**B** Left). On the other hand, SKAT(Avg) and mKU identify a large number of modules (Figure 6**B** Right), which are consistent to the inflated type 1 errors of these two methods as observed previously, and failed to provide specific prioritizations of the gene modules. Remarkably, by comparing to the significant GWAS SNPs identified from another genome-wide meta-analysis of brain volume study (58), two out of the four selected modules by BayesKAT (50%) contain significant GWAS SNPs (Figure 6**C**). In contrast, only 26% (7/27) and 43% (6/14) modules selected by mKU and SKAT(Avg) contain significant GWAS SNPs. These results further provide orthogonal support for the superior performance of BayesKAT in identifying new collective associations to the complex traits for SNP groups of functionally related genes.

When applied to SNP groups organized according to PPI modules that largely represent protein complexes, BayesKAT distinctly prioritizes eight PPI modules as significant, six of which contain significant GWAS SNPs (75%, Figure 6**D**, Figure 6**E** Left). On the contrary, only 30%(17/56) and 65%(13/20) of the modules selected by mKU and SKAT(Avg) respectively contain significant GWAS SNPs (Figure 6**D**, Figure 6**E** Right). Taken together, the highly specific prioritizations of potential gene modules and protein complexes, along with the substantially improved justification from other GWAS SNPs, suggest that BayesKAT can facilitate novel discoveries of molecular components involved in complex traits and may pave the way for innovative approaches to disease treatments.

## DISCUSSION AND CONCLUSION

BayesKAT (https://github.com/wangjr03/BayesKAT) is a flexible and data adaptive methodology that automatically selects the appropriate composite kernel using the MCMC algorithm (BayesKAT-MCMC) or the optimization technique (BayesKAT-MAP) and conducts hypothesis testing on the presence of group-level genetic associations for complex traits. The Bayesian framework and the inferred model outcomes of posterior probabilities are more interpretable and informative, compared to the calculated p-values from the frequentist methods. Based on the extensive simulation benchmark analyses, BayesKAT demonstrates consistent and robust superior performance than other methods across different settings. Moreover, evaluated on a series of biologically inspired SNP groups based on a real genetic dataset, BayesKAT not only achieves improved prioritization of functionally relevant and justified group-level SNP associations, but also enables novel discoveries with respect to the underlying molecular mechanisms of complex traits. By revealing the collective effects of functionally cooperative SNPs without relying on the prior knowledge of specific kernels, BayesKAT represents one important step forward towards the goal of deciphering the intricate genetic basis of human diseases.

Although some methods based on the Gaussian process (59) or supervised learning technique (60) attempt to select the best kernel using training data for prediction purposes, BayesKAT is the first Bayesian KBT methodology that simultaneously selects the optimal composite kernel while testing for the associations, without requiring the training data. In addition, the data-adaptive strategy of composite kernel selections also facilitates the description of more complicated interdependence structures of the data that can not be fully captured by individual kernels. Furthermore, BayesKAT provides the flexibility of incorporating multiple testing corrections, integrating prior knowledge based on biological experiments into the model, and modeling various data types (e.g. discrete SNP features and continuous gene expression features). To complement the MCMC strategy, BayesKAT-MAP is highly scalable and can be efficiently implemented for large-scale genome-wide studies.

BayesKAT utilizes the Metropolis-Hastings MCMC algorithm in combination with a derivative-free grid-search-based optimization approach to choose the composite kernel for specific datasets. Nevertheless, the BayesKAT framework is not restricted to these techniques. Other efficient MCMC sampling algorithms or reliable optimization techniques can be incorporated. A variety of sampling techniques (e.g., bridge sampling, importance sampling, path sampling, etc.) have been reviewed and compared for different purposes of modeling (61), (62), (63), (64). Integrating these techniques, especially the variational Bayes techniques, into the BayesKAT framework and systematically evaluating their performance for different types of applications will be an important step for future developments.

In recent years, different deep learning models have been developed and applied in genomics studies to predict molecular features, such as gene expression, histone marks, chromatin accessibility and transcription factor binding, using DNA sequences as features (65), (66), (67), (68). These models allow for in silico mutations of DNA sequences and the prediction of perturbed molecular features for each mutation individually. Compared to these models, the power of BayesKAT in delineating the collective group-level genetic associations based on automatic composite kernel selection opens up a new level of analytical capability of dissecting the genetic complexity. Combined together, the complementary advantages of BayesKAT and deep learning models are expected to facilitate novel mechanistic insights into human diseases.

The implementation of BayesKAT is not limited to genetic studies based on features of SNPs or gene expressions. Any group of continuous or discrete features that are functionally related can be tested for associations with an outcome using the BayesKAT methodology. For instance, BayeskAT can be used to test if a group of images is associated with a particular disease trait by adopting properly defined kernels. As another direction for future developments, non-linear functions of candidate kernels, instead of the linear combinations, will be explored as the composite kernel, which may lead to improved power for kernel-based testing.

## ACKNOWLEDGEMENTS

This work was supported, in part, by awards R01GM131398 from the National Institutes of Health and NSF1942143 from the National Science Foundation. We thank MSU iCER for providing the high-performance computing infrastructure.

Data collection and sharing for this project was funded by the Alzheimer’s Disease Neuroimaging Initiative(ADNI) (National Institutes of Health Grant U01 AG024904) and DOD ADNI (Department of Defense award number W81XWH-12-2-0012). ADNI is funded by the National Institute on Aging, the National Institute of Biomedical Imaging and Bioengineering, and through generous contributions from the following: AbbVie,Alzheimer’s Association; Alzheimer’s Drug Discovery Foundation; Araclon Biotech; BioClinica, Inc.; Biogen; Bristol-Myers Squibb Company; CereSpir, Inc.; Cogstate; Eisai Inc.; Elan Pharmaceuticals, Inc.; Eli Lilly and Company; EuroImmun; F. Hoffmann-La Roche Ltd and its affiliated company Genentech, Inc.; Fujirebio; GE Healthcare; IXICO Ltd.; Janssen Alzheimer Immunotherapy Research & Development, LLC.; Johnson & Johnson Pharmaceutical Research & Development LLC.; Lumosity; Lundbeck; Merck & Co., Inc.; Meso Scale Diagnostics, LLC.; NeuroRx Research; Neurotrack Technologies; Novartis Pharmaceuticals Corporation; Pfizer Inc.; Piramal Imaging; Servier; Takeda Pharmaceutical Company; and Transition Therapeutics. The Canadian Institutes of Health Research is providing funds to support ADNI clinical sites in Canada. Private sector contributions are facilitated by the Foundation for the National Institutes of Health (www.fnih.org). The grantee organization is the Northern California Institute for Research and Education, and the study is coordinated by the Alzheimer’s Therapeutic Research Institute at the University of Southern California. ADNI data are disseminated by the Laboratory for Neuro Imaging at the University of Southern California.

## DATA AVAILABILITY

BayesKAT is an open source infrastructure available in the GitHub repository https://github.com/wangjr03/BayesKAT. In addition to the R codes, the instructions and sample testing data are also provided.

## Footnotes

1 Data used in preparation of this article were obtained from the Alzheimer’s Disease Neuroimaging Initiative (ADNI) database (adni.loni.usc.edu). As such, the investigators within the ADNI contributed to the design and implementation of ADNI and/or provided data but did not participate in analysis or writing of this report. A complete listing of ADNI investigators can be found at: http://adni.loni.usc.edu/wp-content/uploads/how_to_apply/ADNI_Acknowledgement_List.pdf

