## Supplementary material for "BayesKAT: Bayesian Optimal Kernel-based Test for genetic association studies reveals joint genetic effects in complex diseases"

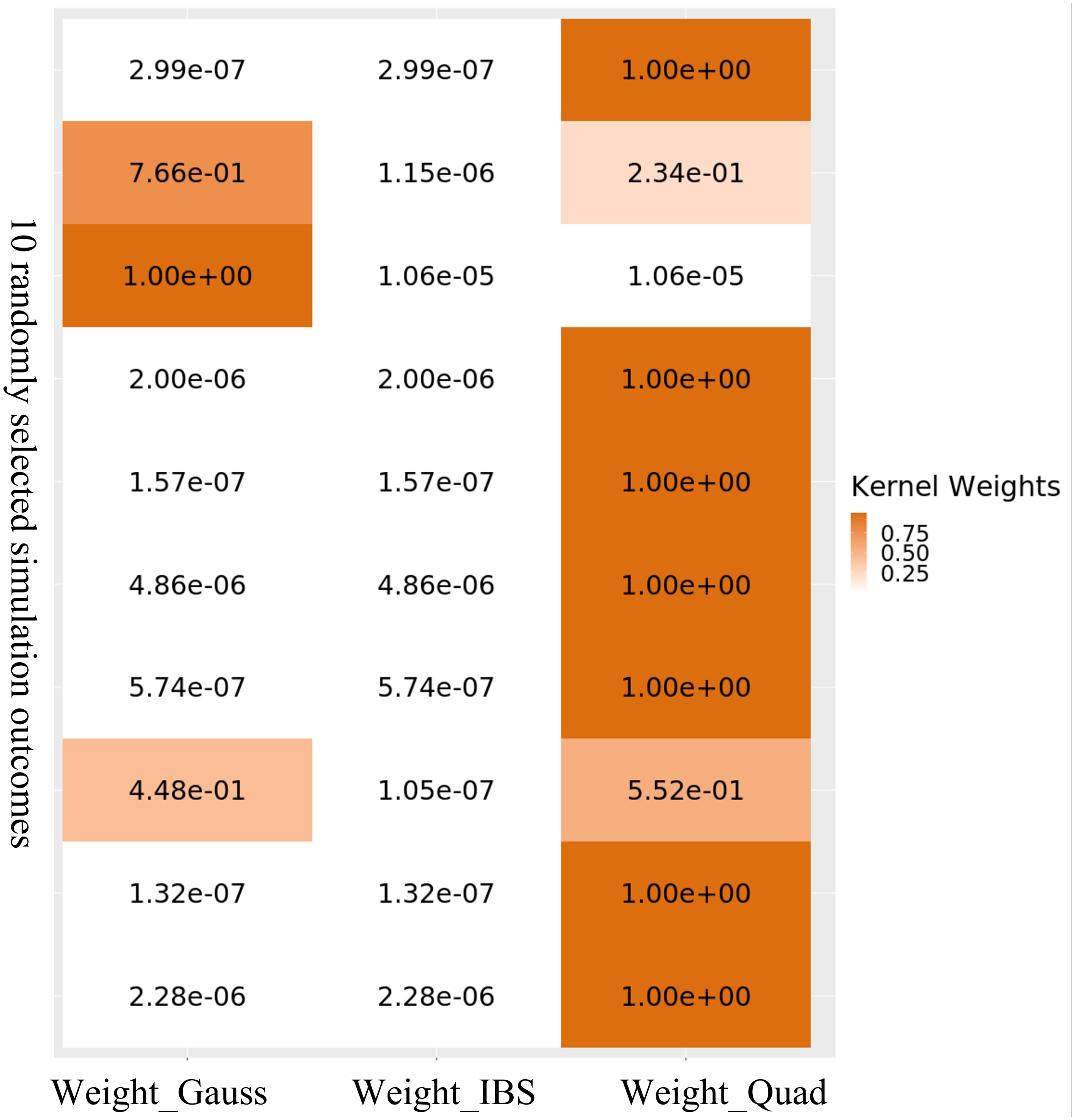


**Supplementary Figure 1**: Inferred weights of each candidate kernel within the composite kernels based on the simulation tests as presented in Figure 1(B). The inferred composite kernel demonstrates strong agreements with the underlying true kernel function (the Quadratic kernel) used for data generation. High kernel weights are consistently inferred for the Quadratic kernel.

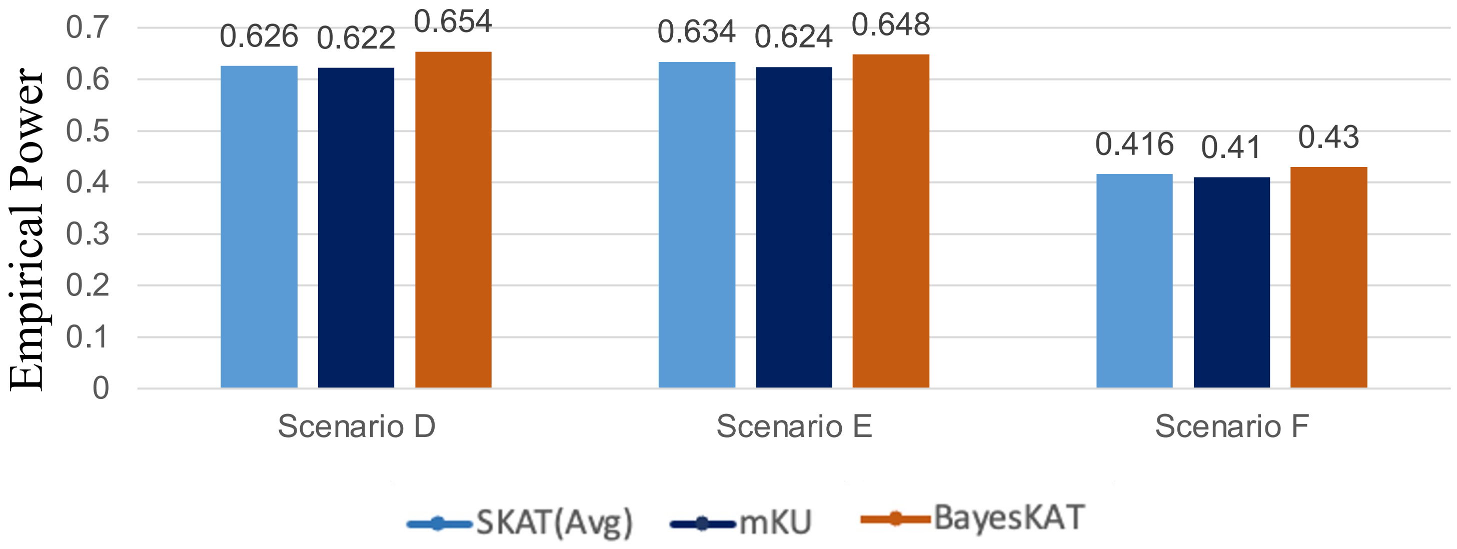


**Supplementary Figure 2**: Performance comparison based on simulations involving unrelated discrete genetic features. With the same fixed level of empirical type 1 error, the empirical power of all three methods are overall low, because this basic simulation only groups unrelated genetic features together, which is not recommended for real-world genetic association studies. Even through, BayesKAT still achieves better empirical power.

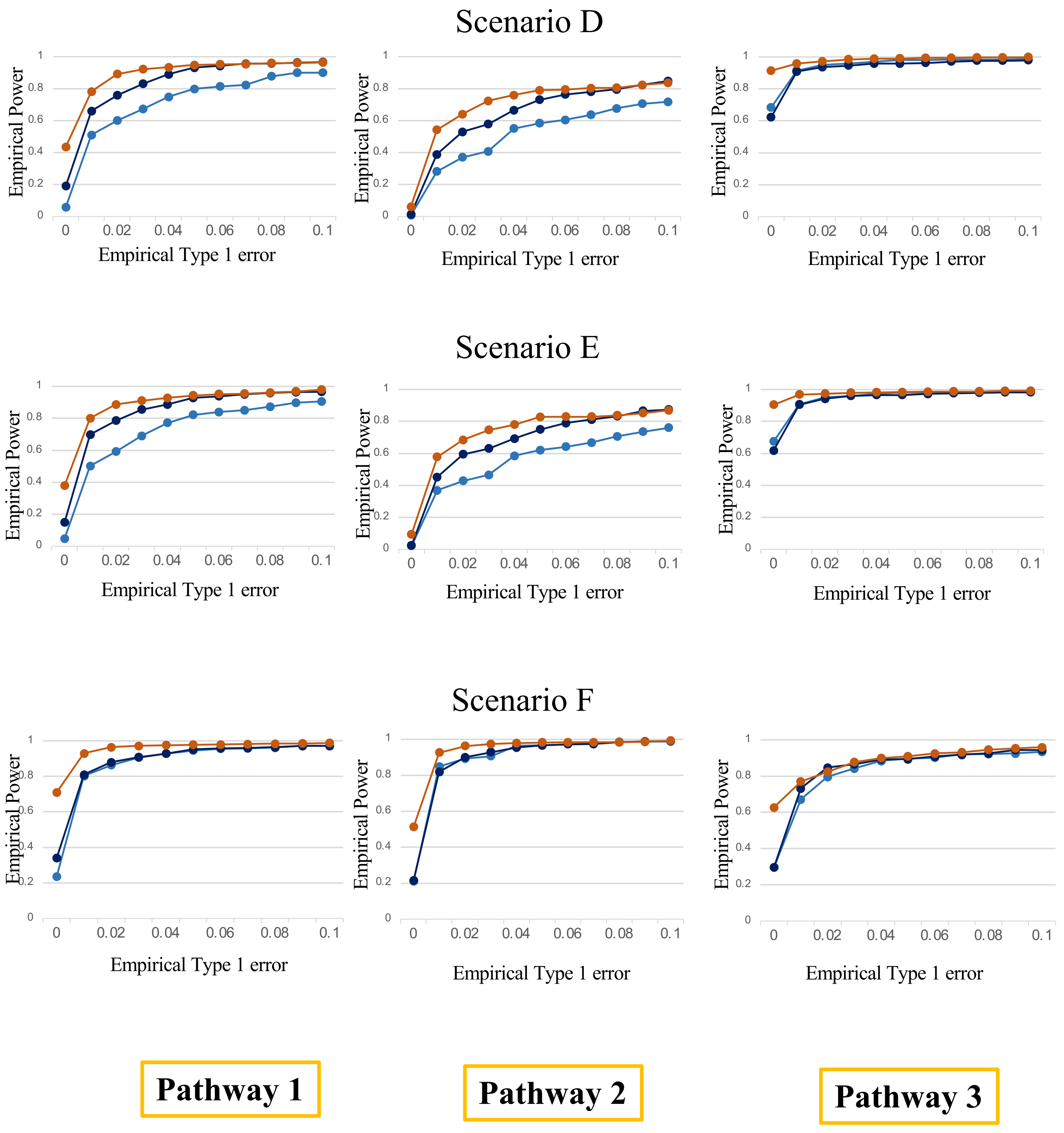


**Supplementary Figure 3**: The empirical power vs empirical type 1 error plot from the pathway-based simulations. For each scenario, BayesKAT consistently achieves the best performance.

| Gene name | P(H1\|Data) |
|---|---|
| 1. CARD10 | 0.9835514 |
| 2. MGAT5 | 0.9474112 |
| 3. TMEM71 | 0.9237384 |
| 4. FAM174B | 0.9042780 |
| 5. ACTA2 | 0.8677653 |
| 6. C18orf45 | 0.8569289 |
| 7. TMEM163 | 0.8357041 |
| 8. LRP1B | 0.8257515 |
| 9. SLC35B4 | 0.8248786 |
| 10. EDNRA | 0.7878737 |
| 11. ABCB7 | 0.7699132 |
| 12. C9orf135 | 0.7660853 |
| 13. SLC25A18 | 0.7583290 |
| 14. DKK2 | 0.7501011 |
| 15. PCID2 | 0.7387756 |
| 16. TMEM38A | 0.7042073 |
| 17. NXNL2 | 0.7037605 |

**Supplementary Table 1**: List of significant genes selected by BayesKAT and their posterior probability of association.

| Pathway name | P(H1\|Data) |
|---|---|
| 1. Pathways of neurodegeneration - multiple diseases | 0.9998840 |
| 2. Alzheimer disease | 0.9982576 |
| 3. Salmonella infection | 0.9897521 |
| 4. Antifolate resistance | 0.9666899 |
| 5. Huntington disease | 0.9630876 |
| 6. Bile secretion | 0.9530779 |
| 7. Alcoholic liver disease | 0.9285561 |
| 8. Dilated cardiomyopathy | 0.9240475 |
| 9. Metabolic pathways | 0.9211165 |
| 10. Amyotrophic lateral sclerosis | 0.9070785 |
| 11. Hypertrophic cardiomyopathy | 0.8975824 |
| 12. Cardiac muscle contraction | 0.8966726 |
| 13. mTOR signaling pathway | 0.8869947 |
| 14. cAMP signaling pathway | 0.8688107 |
| 15. Pathogenic Escherichia coli infection | 0.8645842 |
| 16. Calcium signaling pathway | 0.7962372 |
| 17. ABC transporters | 0.7851350 |
| 18. Citrate cycle (TCA cycle) | 0.7766141 |
| 19. Non-alcoholic fatty liver disease | 0.7676582 |
| 20. AMPK signaling pathway | 0.7640913 |
| 21. Parkinson disease | 0.7242965 |

**Supplementary Table 2**: List of the significant KEGG pathways selected by BayesKAT and their corresponding posterior probabilities of association.
